# Genotypic and resistance profile analysis of two oat crown rust differential sets urge coordination and standardisation

**DOI:** 10.1101/2023.09.28.559455

**Authors:** Duong T. Nguyen, Eva C. Henningsen, David Lewis, Rohit Mago, Meredith McNeil, Radoslaw Suchecki, Scott Boden, Jana Sperschneider, Shahryar F. Kianian, Peter N. Dodds, Melania Figueroa

## Abstract

*Puccinia coronata* f. sp. *avenae* (*Pca*) is the causal agent of the disease known as crown rust which represents a bottleneck in oat production worldwide. Characterisation of pathogen populations often involves race (pathotype) assignments using differential sets, which are not uniform across countries. This study compared virulence profiles of 25 *Pca* isolates from Australia using two host differential sets, one from Australia and one from the USA. These differential sets were also genotyped using DArT sequencing technology. Phenotypic and genotypic discrepancies were detected on eight out of 29 common lines between the two sets, indicating that pathogen race assignments based on those lines are not comparable. To further investigate molecular markers that could assist in the stacking of rust resistance genes important for Australia, four published *Pc91*-linked markers were validated across the differential sets and then screened across a collection of 150 oat cultivars. Drover, Aladdin, and Volta were identified as putative carriers of the *Pc91* locus. This is the first report to confirm that the cultivar ‘Volta’ carries *Pc91* and demonstrates the value of implementing molecular markers to characterise materials in breeding pools of oat. Overall, our findings highlight the necessity of examining seed stocks using pedigree and molecular markers to ensure seed uniformity and bring robustness to surveillance methodologies.

## Introduction

Oat (*Avena sativa* L.) is a grain crop that is in high demand due to its desirable nutritional properties for humans and animals, as well as for various commercial applications such as cosmetics and fibre production (Rasane et al. 2015; Suttie and Reynolds 2004). Australia needs to increase oat production to meet the growing demands from domestic and international markets, especially in Asia and Africa (Agriculture and Food 2022; Cowman et al. 2021). Crown rust disease, caused by the rust fungus *Puccinia coronata* f. sp. *avenae* (*Pca*), severely impacts oat production worldwide (Nazareno et al. 2018; Simons 1985). Crown rust may cause yields to drop by 10% to 40%, and under optimal environmental conditions, *Pca* infections can destroy entire fields (Behnken et al. 2011; Carson 2017; Strunk et al. 2022). In Australia, the disease can result in substantial yield losses of up to 50% in susceptible varieties (Chambers and Thomas 2020).

Oat crown rust disease is managed via two methods, chemical control and planting genetically resistant cultivars (Barr 1994). Genetic manipulation is the preferred method to prevent rust epidemics as it minimises adverse effects of agrochemicals to the environment, harmful residues in the crop, and costs associated with treatments (Figueroa et al. 2020). Resistance manifests either as broad spectrum (e.g., partial or adult plant resistance) or race-specific (Periyannan et al. 2017). Race-specific resistance, also known as seedling or all-stage resistance (ASR), is advantageous because it is effective from seedling growth stages onwards and provides full immunity in some cases, while adult plant resistance (APR) can be more durable. In the best-case scenario, a combination of both types of resistance is utilised to achieve the best outcomes (Ellis et al. 2014). The underlying mechanism controlling race-specific resistance involves the recognition of pathogen-derived proteins, known as avirulence (*Avr*) effectors, by plant immune receptors encoded by resistance (*R*) genes, which trigger a plant defence response and stop pathogen infection. However, the ability of pathogens to evolve variants of *Avr* effectors and avoid plant recognition poses a limitation to this strategy (Figueroa et al. 2020). The evolution of *Pca* can also be driven by sexual processes, which contribute to generating new virulence traits through the reassortment of virulence alleles (Miller et al. 2020; Nazareno et al. 2018). Common buckthorn (*Rhamnus cathartica*) is the sexual host of *Pca* and the lack of this species in Australia has led researchers to postulate that *Pca* evolution relies solely on asexual propagation in this part of the world (Harder et al. 1992; Simons 1985).

As of today, 92 loci (*Pc1-Pc85, Pc91-Pc96, Pc98*) associated with *Pca* resistance are nominated based on phenotypic data and limited genetic characterisation (Carson 2017; Nazareno et al. 2018). Some of these have been genetically mapped in oats (Carson 2017; Park et al. 2022a). Furthermore, none of the *Pc* genes have been cloned so far, and only a handful of tightly linked molecular markers have been developed. This makes it difficult to assess whether the same *R* genes or alleles of the same *R* gene have been independently identified and redeployed under different names. Unfortunately, reports from the global oat community indicate that most of these genes are now ineffective in many regions as *Pca* populations have evolved to overcome their resistance. A potential strategy to deliver rust-durable resistance is the stacking of race-specific and broad-spectrum resistance genes (Ellis et al. 2014). Such an approach requires the use of molecular markers to select for each *R* gene in germplasm and effectively incorporate multiple different sources of resistance into the same cultivar (Ellis et al. 2007; Revathi et al. 2010)

At present, rust surveillance programs conducted by pathologists rely on race assignments (pathotype) to monitor rust evolution and movement and inform breeding programs and growers about which oat cultivars are suitable for seasonal planting. Races of rust fungi are commonly defined by a standard nomenclature system that links virulence phenotypes to race-specific genes postulated in subsets of crop lines, referred to as differential sets (Chong et al. 2000; Figueroa et al. 2018; Leonard 2016). This strategy, although limited to a small set of markers (*R* genes) has provided important insights into the fast-evolving nature of crown rust populations regardless of geographic origin (Carson 2011). While several *Pc* genes and oat differential sets are shared among international research organisations, there is no universal collection due to regional differences in pathogen virulence and host resistance (Chong et al. 2000). The oat differential set used in Australia includes up to 47 lines, 29 of which overlap with lines included in the United States of America (USA) oat differential set of 40 lines (Nazareno et al. 2018). The oat lines in the differential set are not isogenic and race assignment reports suggest that several oat differential lines may carry multiple *Pc* genes (Carson 2017; Park 2013), and some *Pc* genes may be duplicated within sets (Hewitt et al. 2023; Miller et al. 2020).

*Pc91* is one of the most recently described race-specific *Pc* genes and was originally identified in the highly crown rust-resistant oat line Amagalon. Amagalon was derived from a cross between tetraploid *A. magna* and diploid *A. longiglumus*, where *A. magna* was the source of disease resistance (Rooney et al. 1994). McCartney et al. (2011) mapped *Pc91* in a recombinant Inbred line (RIL) population (n=100) genotyped using Diversity Array Technology (DArT) and found several co-segregating DArT markers. Subsequently, Gnanesh et al. (2013) used the sequence of DArT marker oPt-0350 (McCartney et al. 2011) to design three co-dominant KASP markers based on three separate SNPs within this sequence (*oPt-0350-KOM4c2*, *oPt-0350-KOM5c1*, and *oPt-0350-KOM6c2*). These markers accurately distinguished oat lines with and without *Pc91* resistance locus in 16 North American oat breeding lines, and oPt-0350-KOM4c2 co-segregated with *Pc91* in three mapping populations and was separated by 1.9cM in a fourth population (Gnanesh et al. 2013). In another study, Klos et al. (2017) mapped *Pc91* to the linkage group Mrg18 through a Genome-Wide Association Study and identified a *Pc91*-associated SNP marker, *GMI_ES03_c2277_336*. Mrg18 corresponds to chromosome 1A in the reference genome assembly *Avena sativa* OT3098 v2 (*Avena sativa* – OT3098 v2, PepsiCo, https://wheat.pw.usda.gov/jb?data=/ggds/oatot3098v2-pepsico), which is the same chromosome where the translocation breakpoint involving then defined chromosome 7C and 17 was located in a Kanota X Ogle map (Tinker et al. 2009). The DArT markers flanking *Pc91* (McCartney et al. 2011, Gnanesh et al. 2013) were aligned with this translocation breakpoint.

In this study, we conducted a genotypic and phenotypic comparison of the widely utilised oat differential sets in Australia and the USA. The primary objective was to pinpoint any discrepancies between lines supposed to represent the same differential in each set. This effort is crucial for enhancing the effectiveness of pathogen surveillance and enabling more precise international comparisons of pathogen race information across countries. Furthermore, given that *Pc91* remains effective in providing crown rust resistance in Western Australia (Henningsen et al. 2023), we also tested the aforementioned *Pc91* molecular markers (Gnanesh et al. 2013; Klos et al. 2017) on a collection of oat cultivars. This aimed to confirm markers’ specificity and identified putative carriers of *Pc91* as a resource for breeding programs. Such information would facilitate marker-assisted selection for durable resistance to crown rust.

## Materials and methods

### Plant material

Two oat crown rust differential sets commonly used in North America (n= 40) and Australia (n=47), respectively, and a collection of oat accessions and cultivars (n=150) were utilised in this study (**Table S1**). The North American differential set was provided from the US Department of Agriculture-Agricultural Research Service (USDA-ARS) Cereal Disease Laboratory (St. Paul, MN, USA) (Nazareno et al. 2018), and lines of the Australian oat differential set (Cuddy et al. 2016; Park 2013; Park et al. 2022b) were accessed from the Australian Grains Genebank (AGG), the Department of Agriculture and Fisheries (DAF), QLD, Australia, and an *Avena* seed stock at CSIRO. We also built an oat collection that includes cultivars with diverse pedigree information by accessing AGG, DAF, and CSIRO-*Avena* resources. These oat lines were amplified through a single seed descent process. Oats were planted at a depth of 1 cm in an in-house soil blend and grown in a glasshouse with a daily cycle of 16 hours at 23°C light and 8 hours at 18°C dark.

### Rust infection assays

A set of 25 *Pca* isolates from major Australian oat-grown regions in 2022 was used (**Table S2,** Henningsen et al. 2023). Infection methods were adapted from Miller et al. (2020), except that rust spores were mixed with talc powder instead of suspended in oil. *Pca* samples were applied at 10 days after sowing by dusting the inoculum onto plants that were pre-misted with water. Plants were kept at high humidity conditions for two days in a sealed inoculation chamber followed by a further 8 days in a growth cabinet (18-23°C, 16h photoperiod). Infection scores were recorded at 10 days post inoculation using a disease rating scale from Murphy (1935) and Nazareno et al. (2018). The scores were subsequently converted to a 0-4 numeric scale for visualisation in heatmaps as described by Miller et al. (2020) and Henningsen et al. (2023), using ComplexHeatmap (v2.14.0) in R (4.2.2) (Gu, 2022) and comparison of virulent impact using violin plots (ggplot2 package) and Wilcoxon rank sum tests in R v4.2.2.

### Genotyping of oat lines and population structure analysis

DNA was extracted from 3-week-old seedlings as described by Ellis et al. (2005) and quality and concentration were assessed using a NanoDrop™ 8000 Spectrophotometer (NanoDrop Technologies Inc., Santa Clara, CA, United States) and fluorescence assay PicoGreen (Singer et al. 1997). DArTSeq genotyping of oat lines was performed by Diversity Arrays Technology Pty Ltd (https://www.diversityarrays.com/) (Canberra, Australia) using restriction enzymes *PstI* and *HpaII* with markers generated in two formats: SilicoDArT (presence/absence variants) and single-nucleotide polymorphisms (SNPs).

Sequences of DArT markers were searched against the reference genome sequence *Avena sativa* OT3098 v2 using BLAST (Altschul et al. 1990) with an expected value (E) < 5e-7 and sequence identity of > 70%. Only markers that matched a genomic location were kept. SilicoDArT and SNP markers were filtered separately using parameters: missing call ≥ 0.5, minimum allele frequency (MAF) < 0.05, and Pearson’s correlation values r > 0.95 (Gezan et al. 2022). After that, SilicoDArT and SNP markers were combined for genotypic analysis, and markers with high correlation (Pearson’s correlation values r > 0.95) were removed.

CMplot (Yin et al. 2021) was used to visualise the genome-wide distribution of DArTSeq markers across oat chromosomes of the OT3098 v2 reference genome. The ASRgenomics package (Gezan et al. 2022) was used in Rstudio (v2022.12.0) to generate a Kinship matrix (G) from DArTSeq genotypic data for Principal Component Analysis (PCA). Phylogenetic analysis was performed using the maximum likelihood criterion. The Python script vcf2phylip.py (Ortiz 2019) (https://github.com/edgardomortiz/vcf2phylip) was used to convert marker vcf files to PHYLIP format for phylogenetic analysis in RAxML (v8.2.12) (Parameters as follows: raxmlHPC-PTHREADS-AVX -f a -# 500 -m GTRCAT -p 12345 -x 12345 -s differential_set.min4.phy -n ML-BS500) (Stamatakis 2014). The output phylogenetic tree was midpoint-rooted and visualised using iTOL (v6.7.2) (Letunic and Bork 2021).

### Kompetitive Allele Specific PCR (KASP) assays for *Pc91*

In this study, we used three KASP markers (*oPt-0350-KOM4c2*, *oPt-0350-KOM5c1*, and *oPt-0350-KOM6c2*, that were reported by Gnanesh et al. (2013). In addition, we converted the GBS-SNP, *GMI_ES03_c2277_336* (Klos et al. 2017) into KASP maker by using the flanking sequence of the SNP to design primers through 3CR Bioscience Free Assay Design Service (https://3crbio.com/). This newly designed KASP marker was named KASP_GMI_ES03_c2277_336, and the sequence of the KASP primers can be found in **Table S3**. The physical locations of these markers in the OT3098 v2 genome reference were determined by completing a BLAST search of the flanking sequence of the marker *GMI_ES03_c2277_336* and the DArT clone oPt-0350 sequence against the reference genome OT3098 v2. The KASP assays were performed in the SNPline PCR Genotyping System (LGC, Middlesex, United Kingdom), using a method previously described by Shi et al. (2023).

### Pedigree visualisation of oat lines

Pedigree records were obtained from the “Pedigrees of Oat Lines” POOL database (Tinker and Deyl 2005; https://triticeaetoolbox.org/POOL/) and from Fitzsimmons et al. (1983). The format was resolved to Parent 1 and Parent 2 to make them compatible with the Helium pedigree viewing software (Shaw et al. 2014) which was used to visualise relationships between accessions. The information of postulated *Pc* genes in oat lines (Cuddy et al. 2016; Park 2013; Tinker and Deyl 2005; https://triticeaetoolbox.org/POOL/) was also imported into Helium for pedigree examination.

## Results

### Comparison of disease resistance phenotypes in oat lines across two host differential sets

We compared the resistance profiles of 87 oat lines (genotypes) (**Table S1**) included in crown rust differential sets from USA and Australia using 25 Australian *Pca* isolates (**Figure 1**). Of the 29 oat accessions that overlap between the two oat differential sets, 13 accessions carrying *Pc* genes (Pc35, Pc38, Pc45, Pc50, Pc52, Pc54, Pc55, Pc56, Pc59, Pc63, Pc64, Pc68, and WIX4361-9) showed strong agreement in infection scores with all 25 *Pca* isolates (**Figure 2**), with only 0-2 isolates detecting a different outcome, resistant (infection type 0 to 2) versus susceptible (infection type 3-4). These small variations could be explained by the biases in visual scoring, and infection variations introduced by factors such as inoculum load and other environmental conditions (e.g., temperature, humidity). However, we detected more substantial phenotypic discrepancies in other lines that should represent the same genotype in each panel. Among these, seven lines (Pc36, Pc39, Pc48, Pc51, Pc58, Pc62, and Pc71) differed in response to 3-4 *Pca* isolates, while nine lines (H548, Pc14, Pc40, Pc46, Pc57, Pc60, Pc61, Pc67, Pc70) showed differences in more than 4 isolates, with some displaying entirely contrasting results (e.g., H548, Pc14, Pc60, Pc70).

**Figure 1:**
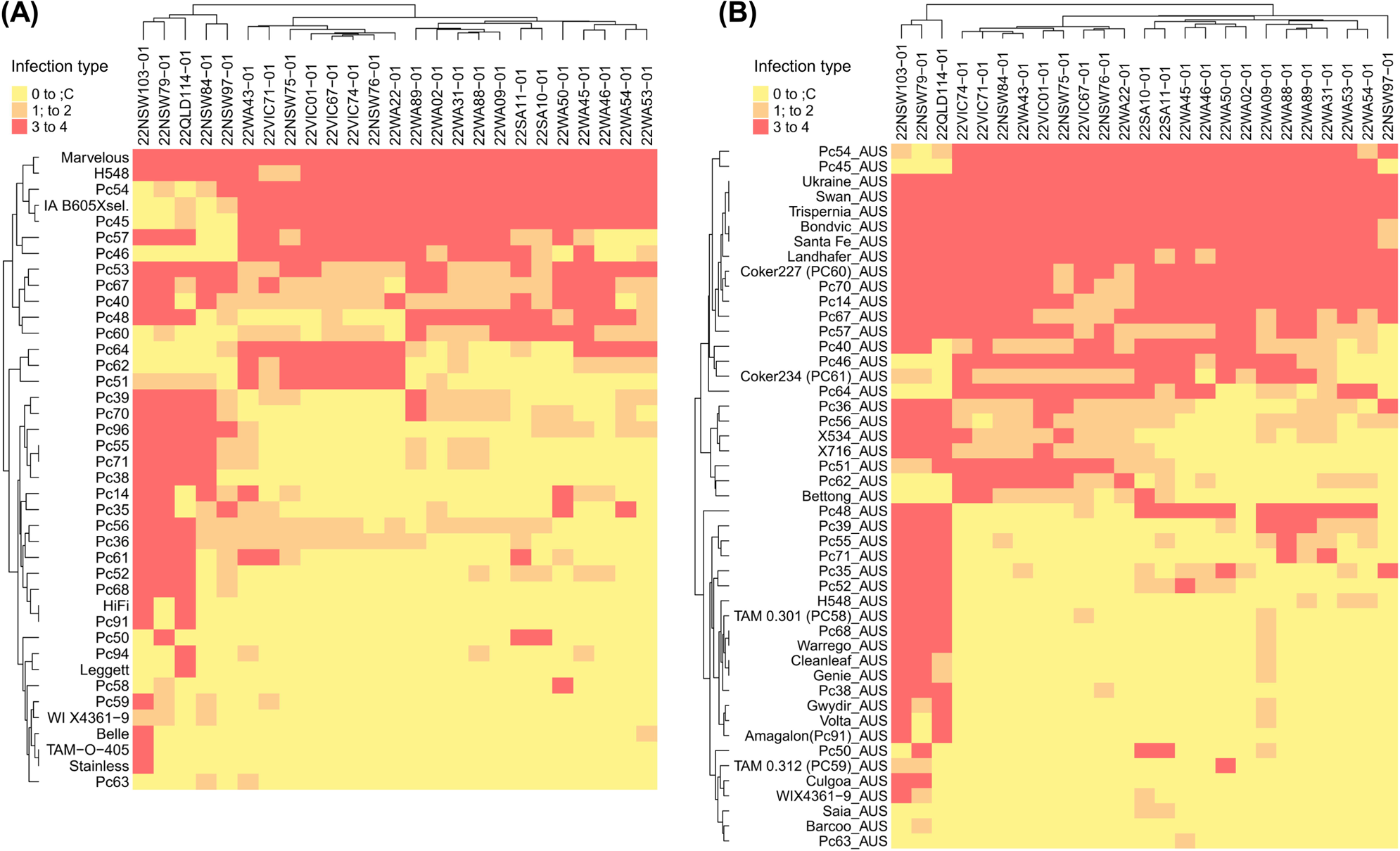
Heatmaps showing the virulence profile of 25 Australian *Pca* isolates. Lower infection types (resistance) are shown in yellow (0 to ;C) and orange (1; to 2), while high infection types (susceptibility) are shown in red (3 to 4). Heatmap using the **(A)** USA oat differential set and **(B)** Australian oat differential set. Dendrograms reflect hierarchical clustering of columns and isolates with similar virulence patterns.

**Figure 2:**
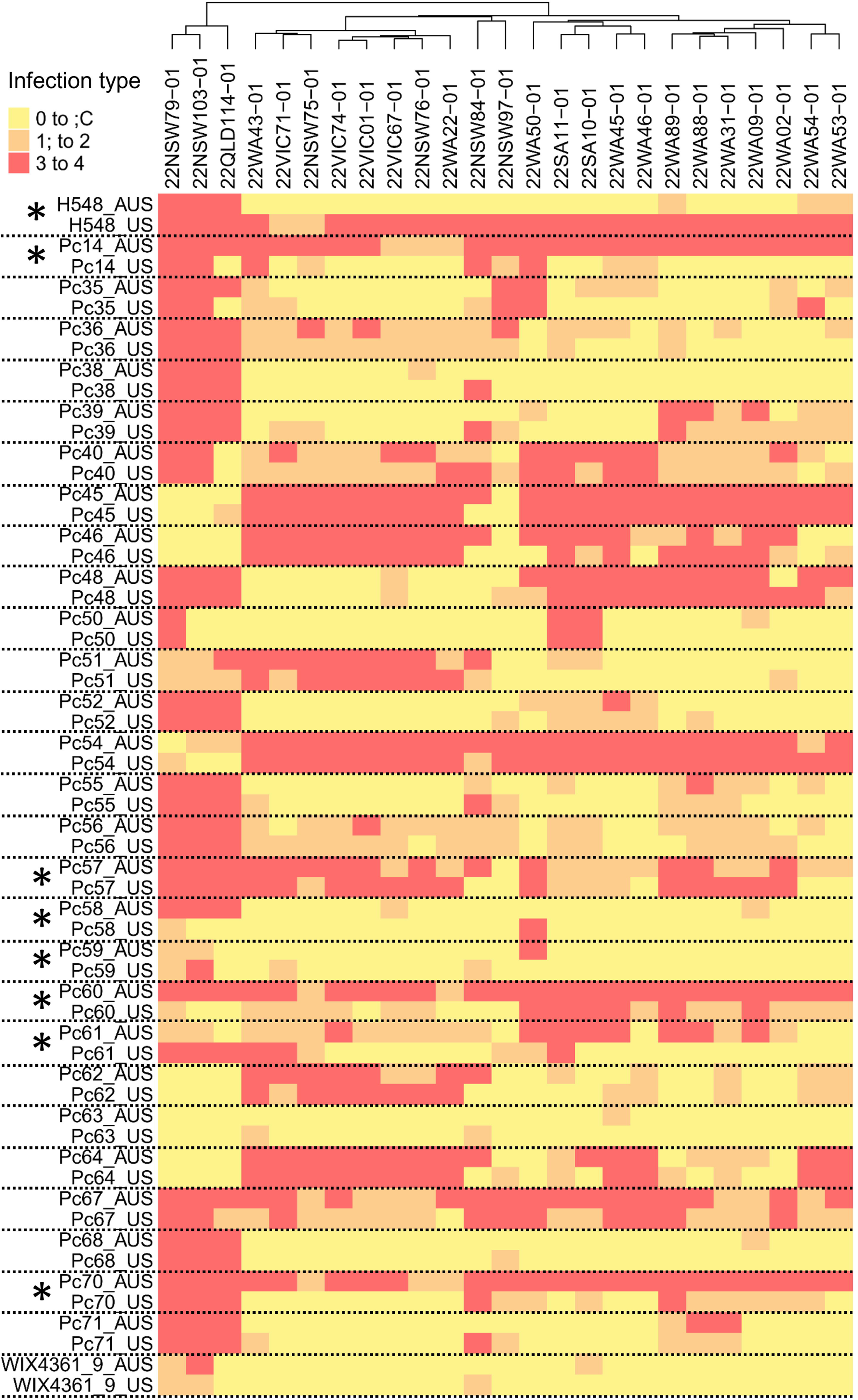
Heatmap shows comparison of virulence profiles of 29 shared differential lines in the Australian and USA sets. Colour range indicates infection type of isolates (columns) on the host lines (rows): low virulence (resistance) in yellow and orange (0 to ;C and 1; to 2) and high virulence (susceptibility) in red (3 to 4). Asterisks indicate pairs that were found to be genotypically divergent in Figure 4. Dendrograms reflect the hierarchical clustering of columns and isolates with similar virulence patterns.

As reported earlier (Henningsen et al. 2023), a higher virulence frequency was observed in the *Pca* isolates from Eastern Australia (QLD, NSW, VIC, and SA) than in those from Western Australia (WA) when tested on oat differential lines from both the USA (*p* = 0.00059) and Australia (*p* = 0.00088) (**Figure S1**). Oat cultivars Barcoo, Saia, and the differential line Pc63 exhibited resistance to all 25 *Pca* isolates, whereas certain oat lines, such as Swan, Ukraine, Marvelous, Santa Fe, Trispernia, Landhafer, Bondvic, Pc45, Pc54, and Pc67 displayed susceptibility to the majority or all isolates. Forty oat lines across both differential sets exhibited resistance to all crown rust isolates from WA but were susceptible to one or more isolates from eastern Australia. These lines were Pc36, Pc38, Pc50, Pc52, Pc55, Pc56, Pc52, Pc58, Pc59, Pc63, Pc68, Pc71, Pc91 (ND894904), Pc94, Pc96, Amagalon, Barcoo, Belle, Bettong, Culgoa, Cleanleaf, Genie, Gwydir, H548, HiFi, Legget, Saia, Stainless, TAM-O-0405, Volta, Warrego, X534, and WIX4361-9. Among these, virulence to Pc91, Pc94, and Pc96, as well as Amagalon (carrying *Pc91*) and HiFi (also carrying *Pc91*) has already appeared in NSW and/or QLD, despite containing recently designated *Pc* genes.

### Genotypic comparison of oat lines included in two differential sets

To examine the genetic relatedness of oat lines in the two differential sets, genotypes were defined using DArTSeq, a genome complexity reduction-based sequencing technology. After applying quality filtering procedures, a total of 48,111 SilicoDArT markers and 17,223 SNPs were retained. Distribution of markers along chromosomes of the oat reference genome OT3098 v2 (**Figure 3)** indicated complete coverage across the genome with the number of SNPs on each chromosome ranging from 1,736 (chromosome 3A) to 4,754 (chromosome 4D). The SilicoDArT and SNP markers were then combined, and highly correlated markers were removed leaving a set of 55,097 markers that was used to examine the genetic relationships between oat lines in the differential sets. A principal component analysis (**Figure S2**) showed one group of oat lines in both the USA and Australian sets were closely related and different from the remainder of the lines which showed no other obvious groupings. This closely related group is composed of a set of near-isogenic lines (NILs) derived from backcrosses with the oat cultivar Pendek for various *Pc* genes (*Pc35*, *Pc38*, *Pc39*, *Pc40*, *Pc45*, *Pc46*, *Pc48*, *Pc50*, *Pc54*, *Pc55*, and *Pc56*) (Carson, 2017). The differential lines Pc61 (USA) and Pc70 (USA) appeared closely related to the Pendek NIL group despite their pedigree not indicating that they are part of NIL-derived sets (Carson 2017). A maximum likelihood (ML) phylogenetic tree (**Figure 4**) supported the close relationships of the Pendek NIL series including Pc61 (USA) and Pc70 (USA). The differential line Pc70 (USA) showed an identical resistance profile to Pc39 (USA) for 25 *Pca* isolates (**Figure 2**), suggesting that it may in fact be a repeat of the Pc39 differential in this study. In addition, two small groups of NILs in either Fraser (Pc62 and Pc63) or Makuru/SunII (Pc64, Pc67, and Pc68) (Carson 2017) were also closely related as expected (**Figure 4**).

**Figure 3.**
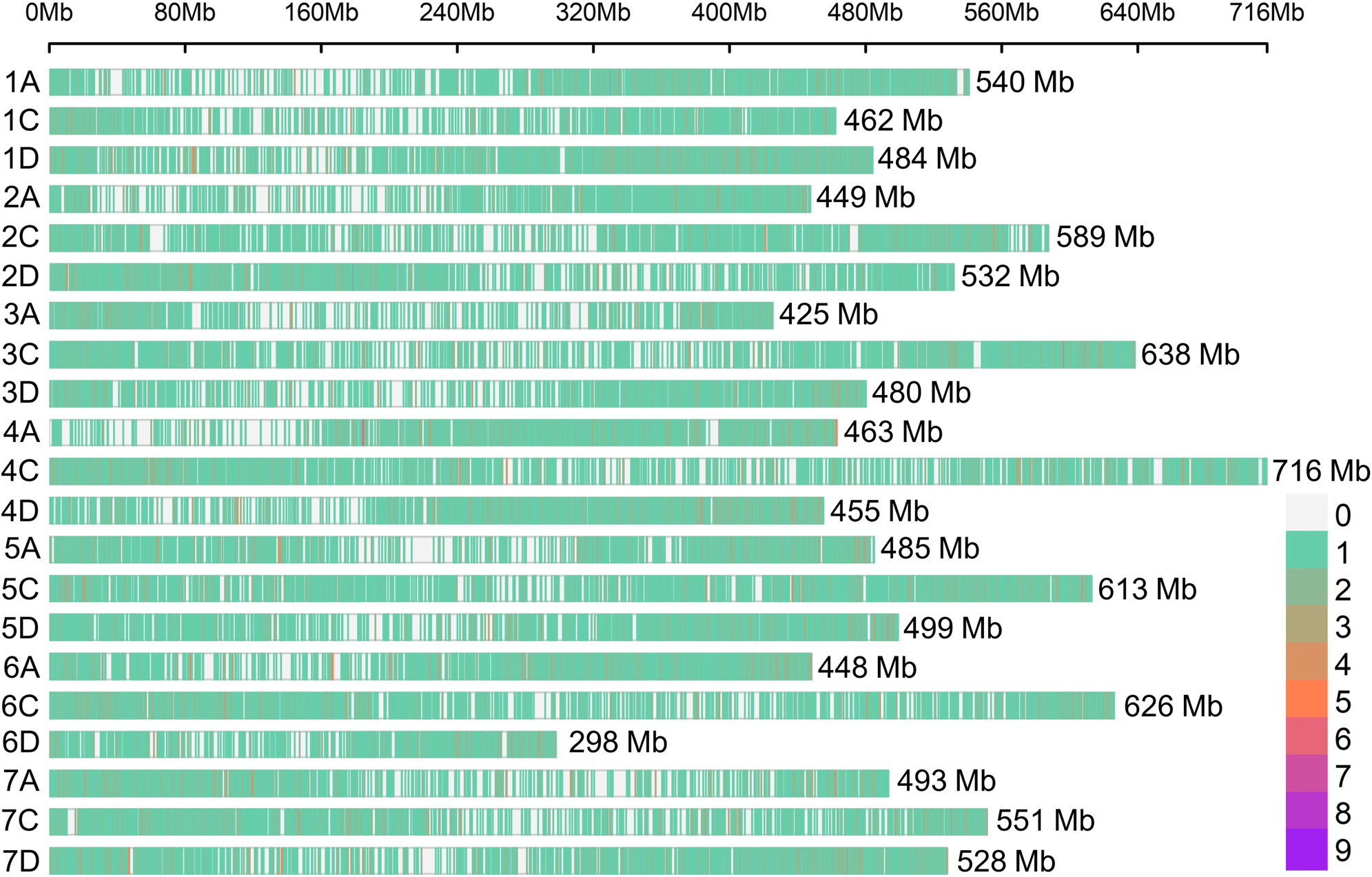
Genome-wide distribution of DArTSeq markers (SNP and SilicoDArT: 65,334) across oat chromosomes of OT3098 v2 reference genome. The lengths of chromosomes are on the right of the coloured bars. The colour scale represents marker numbers within a 1Kb window size.

**Figure 4.**
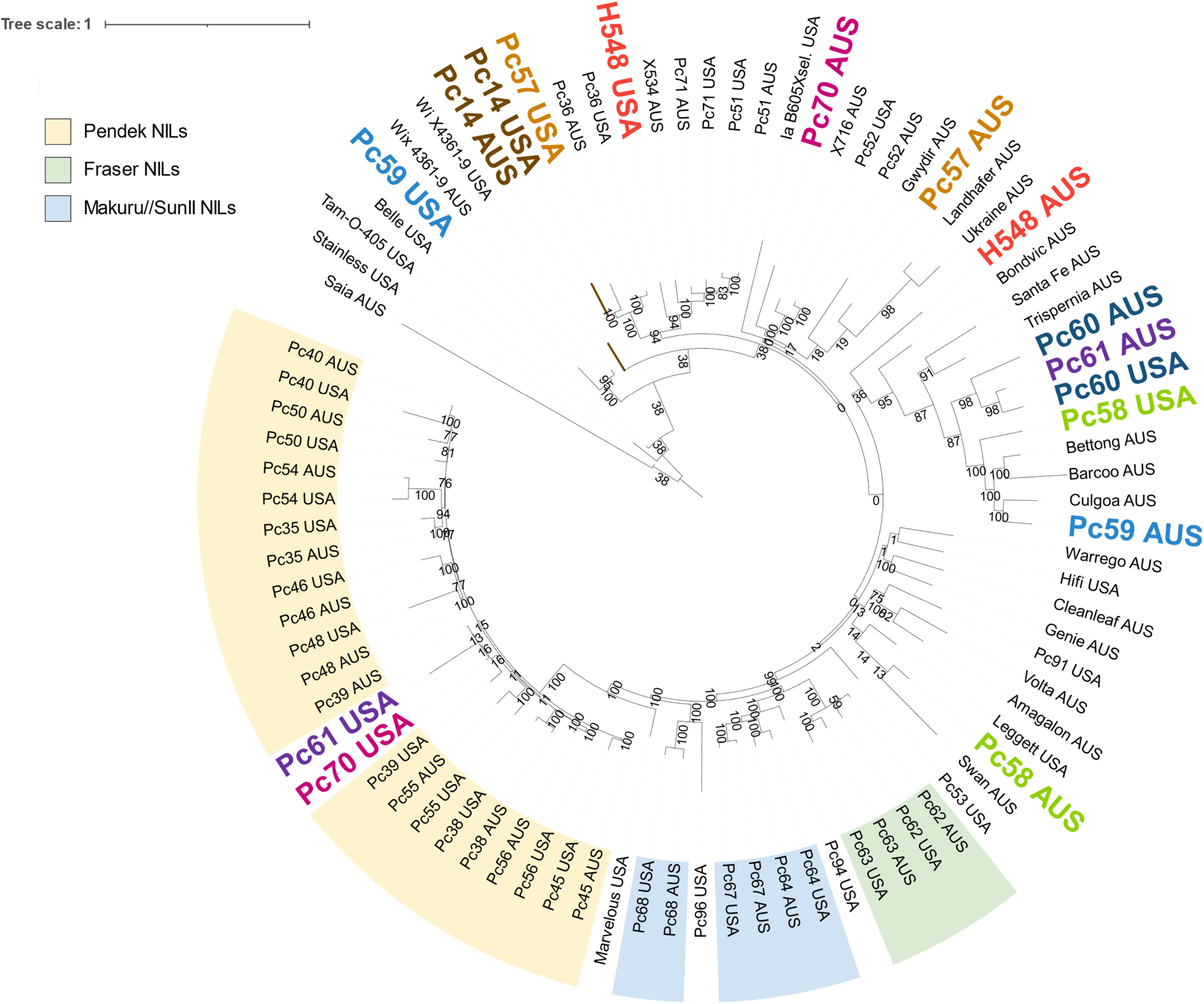
Phylogenetic tree of lines/cultivars present in the USA and Australian oat differential set. The phylogeny was generated based on 55,097 markers (SNPs and SilicoDArT) called against the reference genome OT3098 v2 with support values from 500 bootstraps shown near branches. Oat lines were sourced mostly from AGG in Australia (AUS) and the USDA-ARS Cereal Disease Laboratory (USA). The same font colour indicates lines that should be the same but showed genotypic discrepancies. Colour shading indicates groups of near-isogenic lines (NILs) derived from backcrossing sources or resistance with oat cultivar Fraser (green), Pendek (yellow), Makuru/SunII (blue).

Importantly, most of the 29 lines common to both Australian and USA sets showed a close phylogenetic relationship, except for eight lines (H548, Pc14, Pc57, Pc58, Pc59, Pc60, Pc61, Pc70). Seven of these also exhibited substantial differences in their virulence profiles against the set of 25 *Pca* isolates (**Figure 2**), while the eighth line (Pc59) appeared to be highly similar for resistance phenotype in both sets. This suggests that while the lines might not be genotypically identical, they still carry the same *R* gene.

The USA and Australian versions of the Pc40 and Pc46 lines also differed substantially in resistance profiles, but both of these differentials are part of the closely related Pendek NIL series which makes it difficult to determine from the phylogeny whether they carry the same R gene or not.

### Implementation of diagnostic *Pc91*-linked KASP markers for screening oat collections

To determine the possible presence of the *Pc91* gene in oat collections of importance to breeding programs in Australia, four markers previously associated with *Pc91* (Gnanesh et al. 2013; Klos et al. 2017) were tested in the oat differential lines. The markers *oPt-0350-KOM4c2*, *oPt-0350-KOM5c1*, and *oPt-0350-KOM6c2* are all derived from the DArT marker oPt-0350 (Gnanesh et al. 2013), and were found physically located at 419,792,088 bp; 419,808,321 bp; and 419,808,388 bp, respectively, on chromosome 1A of the reference genome OT3098 v2 PepsiCo, while the SNP marker *GMI_ES03_c2277_336* (Klos et al. 2017) occurs at 387,531,571 bp on the same chromosome (data not shown). The marker *KASP_GMI_ES03_c2277_336* was designed based on the *GMI_ES03_c2277_336* flanking sequence. For breeding purposes, Amagalon is commonly used as the donor source of *Pc91*. Both ND94904 and HiFi genotypes were derived from Amagalon (**Figure 5**) and previously identified as *Pc91* carriers (Gnanesh et al. 2013; McCartney et al. 2011; Rooney et al. 1994). We therefore considered these as positive controls to validate KASP markers of *Pc91* in the oat differential lines.

**Figure 5.**
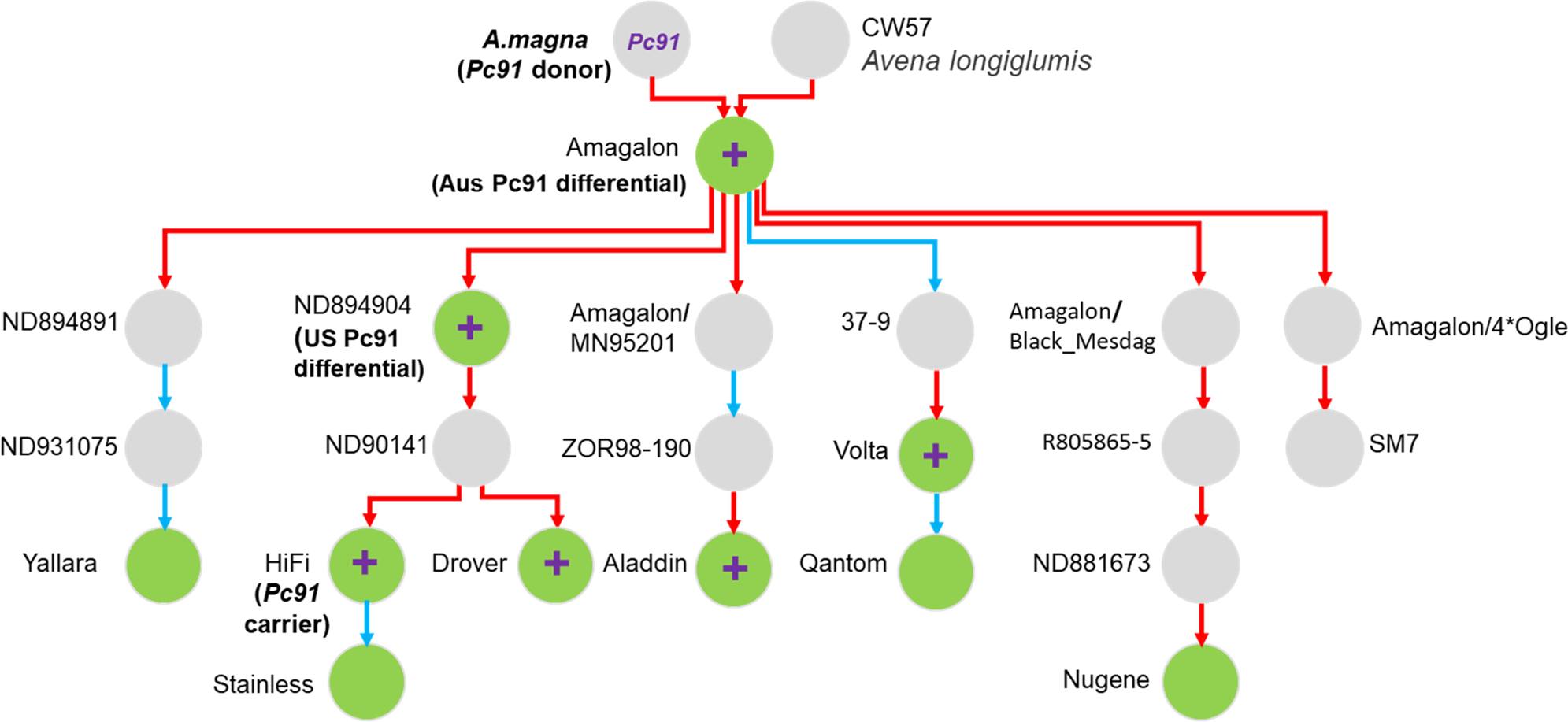
Pedigree of *Pc91* carrier, Amagalon. A simplified pedigree of the cultivar Amagalon was modified from a Helium output (Shaw et al. 2014). Pedigrees were obtained from the “Pedigrees of Oat Lines” POOL database (https://triticeaetoolbox.org/POOL/; Tinker and Deyl, 2005) and Fitzsimmons et al. (1983). Red line indicates maternal parent and blue indicates paternal parent. Green circles indicate lines included in the Differential Sets and oat germplasm collection and tested with *Pc91*-linked markers, with lines positive for these markers indicated by “+”. ND894904, Amagalon, and HiFi were previously defined as *Pc91* carriers (Gnanesh et al. 2013).

KASP assay results with the *oPt-0350-KOM4c2*, *oPt-0350-KOM5c1*, and *oPt-0350-KOM6c2* markers correlated completely (**Table S4**) and detected four lines containing the resistance-associated alleles in the differential sets (n=87). These include the three positive control lines HiFi, which is homozygous for the resistant alleles, and Pc91 (ND894904) and Amagalon, which appeared heterozygous for the resistance and susceptibility alleles **(Figure 6** and **Table S4**). McCartney et al. (2011) also found that Amagalon and the Pc91 differential line (ND894904) carried both alleles of several markers in this region and suggested this was due to a tandem duplication. The line Volta-1 also appeared heterozygous for the markers and exhibited an identical rust resistance profile to HiFi, ND894904, and Amagalon (**Figure S3**). The pedigree of Volta (**Figure 5**) includes Amagalon as a progenitor, so it likely inherited *Pc91* from this source, including the duplicated marker. In contrast to previous reports (Gnanesh et al. 2013; McCartney et al. 2011), the oat cultivar Stainless, which was derived from a cross of ND931475/AC Assiniboia//HiFi (Mitchell Fetch et al. 2011), was not positive for resistance-associated alleles of all tested markers (**Table S4**) and showed different resistance profiles in comparison to other known *Pc91* carriers (HiFi, ND894904, and Amagalon) (**Figure S3**). Moreover, the cultivar Stainless shows significant divergence in the phylogenetic tree from its parental line HiFi (**Figure 4**). This discrepancy raises the possibility that the current Stainless in the USA differential set differs from the source material used in the studies by Gnanesh et al. (2013), and McCartney et al. (2011).

**Figure 6.**
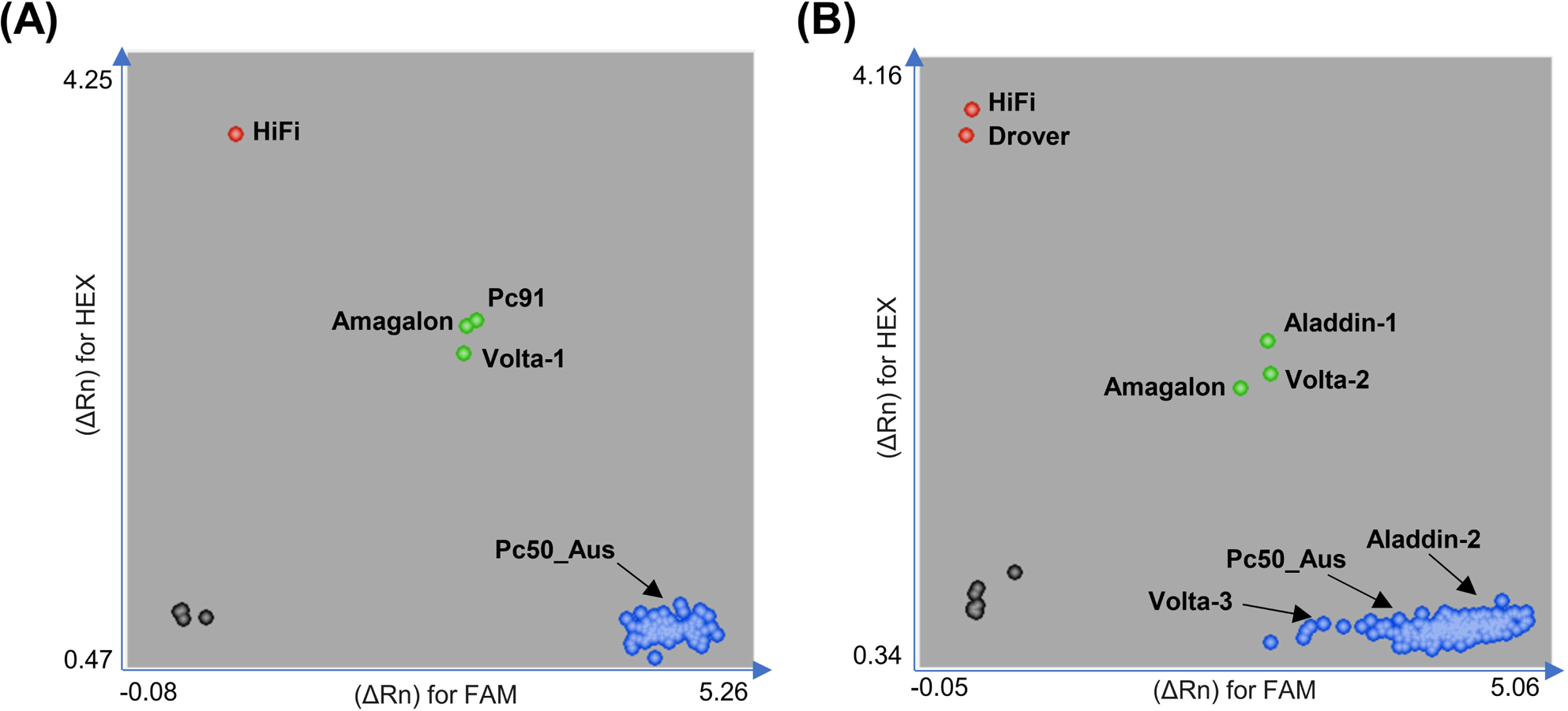
Genotype cluster plots from KASP assays for *Pc91-*linked marker *oPt-0350-KOM4c2* (Gnanesh et al. 2013). **(A)** Differential Sets, n=88. **(B)** Oat Collection, n =150. The markers *oPt-0350-KOM5c1* and *oPt-0350-KOM6c2* displayed a similar pattern as *oPt-0350-KOM4c2*. Blue and red dots show homozygous genotypes for the resistant (*CC*) or susceptible (*TT*) phenotypes, respectively; green dots show heterozygous genotypes (*CT*) for disease resistance; black dots represent non-template control. X-and Y-axes show relative amplification units (ΔRn) for fluorescein amidite (FAM) and Hexachloro-fluorescein (HEX) signals. Aladdin-1, Volta-1 and Volta-2 were derived from AGG seed source (F_6_-F_7_), while Aladdin-2 and Volta-3 were derived from a Breeder’s seed source (F_15+_); DNA from oat lines Amagalon, HiFi, and Pc50_Aus from the differential sets were used as experimental controls in the KASP assays of the oat collection.

The marker *KASP_GMI_ES03_c2277_336* detected six differential lines as positive for the resistance-associated allele, ND94904, HiFi, Cleanleaf, Pc58 (Aus), Warrego, and Leggett, but missed Volta-1 and Amagalon. No pedigree relationship (**Figure 5** and https://oat.triticeaetoolbox.org/) was evident between known *Pc91* carriers (ND94904, Amagalon, and HiFi) and Cleanleaf (carrier of *Pc38*, *Pc39*, *Pc52*), Pc58_Aus, Warrego (carrier of Pc61), and Leggett (carrier of *Pc68*, *Pc94*) (Chong et al. 2000, Mitchell Fetch et al. 2007, Park 2013). In addition, they displayed distinct rust resistance profiles (**Figure S3**). Klos et al. (2017) previously scored the *GMI_ES03_c2277_336* SNP marker on 31 lines of the US differential set, with identical results on these lines to those obtained here using *KASP_GMI_ES03_c2277_336* marker (i.e., only Pc91 was positive).

The *Pc91* markers were further used to screen a collection of 150 oat cultivars (**Table S1**) mostly sourced from AGG with a few exceptions. The oat cultivars Aladdin, Volta, Genie, and Wizard were sourced from both AGG and a DAF breeder, representing different generations of selection. For instance, Aladdin-1 is a single seed stock derived from an AGG seed source representing an F_6_-F_7_ generation of this line, while Aladdin-2 is a single seed stock derived from a breeder’s seed source representing an F_15_+ generation. Similarly, Volta-1 (in the Australian differential set above) and Volta-2 (in the oat cultivar set) are separate single seed stocks derived from AGG (F_6_-F_7_) and Volta-3 is the single seed stock derived from a Breeder’s seed source (F_15_+).

KASP assays with *oPt-0350-KOM4c2*, *oPt-0350-KOM5c1*, and *oPt-0350-KOM6c2* detected only three lines (out of 150) containing the resistance-associated alleles: Drover, Aladdin-1, and Volta-2, with the former homozygous, and the latter two appearing heterozygous (**Figure 6** and **Table S4**). The pedigree visualisation confirmed that these lines have Amagalon or HiFi as parental lines (**Figure 5**). The oat cultivar Drover showed the same rust resistance profile as all the known *Pc91* carriers (HiFi, Amagalon, and ND94904) (**Figure S3**). However, the Aladdin-2 and Volta-3 lines sourced from the breeder were homozygous for the susceptible genotype, indicating genetic differences between the sources. This was supported by a comparison of the resistance profiles of these lines (**Figure S3**). Volta-1 (in the differential set) showed the same resistance profile as Amagalon, Pc91, and HiFi, in contrast, Volta-3 showed a different profile that was identical to Pc50 (USA and Australian sets), while Volta-2 showed a profile consistent with containing both *Pc91* and *Pc50*. The Pc50 differential line was also present in the pedigree of Volta (**Figure S4**). Likewise, the oat cultivars Aladdin-1 and Aladdin-2 showed different resistance profiles, with the former similar to Amagalon, HiFi, and ND94904, but with resistance to one additional isolate (**Figure S3**). Five of our lines (HiFi, AC Assiniboia, Leggett, AC Ronald, and CDC Dancer) were among 16 North American oat breeding lines analysed by Gnanesh et al. (2013) with these markers and gave identical results.

The marker *KASP_GMI_ES03_c2277_336* detected the resistance allele in 27 of the 150 cultivars (**Table S4**). Most of these had never been postulated to carry the *Pc91* locus and only Drover, Aladdin (selections 1 and 2), Volta, and two other oat cultivars, Nugene and Yallara, contained known *Pc91* sources in their pedigree (**Figure 5**, **Figure S4**, and https://oat.triticeaetoolbox.org/). Like the oPt-0350 markers, *KASP_GMI_ES03_c2277_336* also detected differences between the Volta selections, although in this case the resistance-associated allele was detected in Volta-3 but not Volta-2.

## Discussion

Pathogen surveillance programs are key to inform breeders about virulence traits and guide development of disease resistant cultivars. Differential sets to categorise pathogen collections and detect changes in virulence are crucial resources for this purpose. In this study, we compared the performance of two separate oat differential sets used in Australia and the USA. By using phylogenetic and principal component analyses based on DArTSeq genotypic data, we identified substantial genetic differences between oat differential lines in the two sets that are supposed to be identical (e.g., H548, Pc14, Pc57, Pc58, Pc59, Pc60, Pc61, Pc70). The differences between these lines were supported by disease resistance phenotypic data obtained after infection with a set of 25 *Pca* isolates, except in the case of Pc59. Thus, these differential lines cannot be considered equivalent when comparing pathogen race information internationally. Given the goals of our study, we are confident in the robustness of our experimental protocols, mitigating the risk of common issues like seed contamination, mislabelling, and planting errors. However, such errors can explain the observations we make. The observed discrepancies may also be explained by seed stock heterogeneity and the outcrossing nature of oat (Coffman and Wiebe 1930; Derick 1933; Fatunla and Frey 1980; Harrington 1932; Murray et al. 2002), which should be carefully considered when undertaking seed amplification. Given the need for standard global surveys of *Pca* population for effective pathogen movement monitoring, we recommend revisiting the core differential set lines and setting a plan for seed distribution that ensures uniformity. Genotypic data of the differential lines used in this study provide a starting point for researchers to take a similar approach and check for inconsistencies.

The gene *Pc91* remains effective against *Pca* in Western Australia (Cuddy et al. 2016; Park 2013, Henningsen et al. 2023). To prolong its effective lifespan, it is critical to combine *Pc91* with other effective *Pc* genes through strategic breeding efforts. Thus, we tested previously published *Pc91-*associated markers (Gnanesh et al. 2013; Klos et al. 2017) for future use in national breeding programs. In our assays, KASP markers derived from DArTSeq marker oPt-0350 were highly specific to lines known or postulated to contain *Pc91*, detecting resistance-associated alleles in the known carriers Amagalon, Pc91 (ND894904) and HiFi, as well as in Drover and some accessions of Volta and Aladdin, which all have *Pc91* carriers in their pedigree and show similar resistance profiles to the known *Pc91* lines. Drover and Aladdin are grazing oat cultivars and have been previously postulated to carry *Pc91* based on phenotypic data derived from a pathogen differential set (Cuddy et al. 2016; Park 2013). Interestingly, Volta and Aladdin have the duplicate alleles of the oPt-0350 markers from Amagalon, while Drover has the single copy allele of HiFi, which is consistent with the pedigree relationships among these lines (**Figure 5**). The marker *KASP_GMI_ES03_c2277_336,* derived from SNP marker *GMI_ES03_c2277_336*, located approximately 32.2 Mb upstream from oPt-0350 showed lower specificity, with the resistance-associated allele being detected in 6 of the differential lines and 27 of the oat cultivars, most of which do not contain known *Pc91* sources in their pedigree, but not in the *Pc91* source line Amagalon. This suggests that this marker is less tightly linked to the *Pc91* locus, and the allele association has been broken by recombination in some lines. Based on these results, the oPt-0350 markers appear highly diagnostic for *Pc91*.

Importantly, the cultivar Stainless, which originated from the cross ND931475/AC Assiniboia//HiFi (Mitchell Fetch et al. 2011), was not positive for all four *Pc91* tested markers in our differential set, as were the other carriers (ND894904, Amagalon, and HiFi) and was phylogenetically distinct from the parental line HiFi (**Figure 4**). Previously, Stainless was defined as a *Pc91* carrier by Gnaneshs et al. (2013) and McCartney et al. (2011). However, it exhibited different resistance patterns to *Pc91* carriers (ND894904, Amagalon, and HiFi) in our study and in the studies by Miller et al. (2020) and Hewitt et al. (2023). These discrepancies suggest that the Stainless line included in these studies either differs from the ones used by Gnanesh et al. (2013) and McCartney et al. (2011) or has undergone genetic segregation at the *Pc91* locus after distribution.

Furthermore, we found some discrepancies in genotype and phenotype for different seed stocks of cultivars Volta and Aladdin. Two seed stocks of the cultivar Volta independently amplified by single seed descent from the same AGG entry (Volta-1 and Volta-2) were positive for *Pc91*, while a third seed stock derived from material donated by a breeder (Volta-3) was not. While Volta-1 showed a similar rust resistance profile to ND894904, Amagalon, and HiFi, Volta-3 shared the same resistance profile as the oat differential line Pc50 and Volta-2 showed a profile that could be explained by the presence of both *Pc91* and *Pc50*. These results again suggest heterogeneous seed stocks and/or changes as the cultivar was advanced to the F_15_+ generation by the breeder since being deposited to AGG at the F_6-7_ generation stage (B. Winter, personal communication). Notably, Volta was first documented with two rust resistance genes (Plant Variety Journal 2005) and later postulated to carry *Pc50* and *Pc68* based on virulence phenotyping and race assignment (Park 2013). However, pedigree information on the origin of Volta shows ancestors including both *Pc50* and *Pc91* donors, but not *Pc68* (**Figure S4**). Thus, it is likely that Volta material has segregated for *Pc50* and *Pc91*, although it is possible that other genotypes have also been present in other stocks used by Park (2013). The cultivar Aladdin had been proposed to contain *Pc50* and *Pc91* (Park 2013) and our results confirm the presence of *Pc91* in Aladdin-1 but not Aladdin-2, which contains an unknown *Pc* gene whose resistance profile is not consistent with *Pc50*. Again, it is possible that other stocks may have included different genotypes, and these discrepancies interfere with the interpretation of infection data on these lines.

The cultivars Yallara and Nugene, with pedigree linked to *Pc91* carriers, are positive for the resistance allele of the marker *KASP_GMI_ES03_c2277_336*. However, oPt-0350 markers argue against the presence of *Pc91* in these cultivars. Given the physical distance between the oPt-0350 markers and *KASP_GMI_ES03_c2277_336,* it is possible that the association between the sequences recognised by these sets of markers has also been broken by recombination. These cultivars have been postulated to carry *Pc38* and *Pc48*, respectively, based on resistance phenotyping with an array of *Pca* isolates (Cuddy et al. 2016; Park 2013).

In conclusion, the rapid host adaptive evolution displayed by *Pca*, challenges the durability of resistant cultivars once deployed in the field. Therefore, strategic plans to stack genes must be supported by the development and adoption of molecular markers. This study also reveals the problems in tracking and maintaining the resistance sources. It is crucial to select and preserve the purity of resistance sources, with the most effective method also being the use of molecular markers. Standardising oat crown rust differential lines is urged for effective global pathogen monitoring. This requires collaborative efforts within the oat community, seamless communication among research groups, and active engagement of genebanks. Ultimately, a systematic approach that incorporates the use of molecular markers, pedigree analysis, and pathology assays as illustrated in this study is key to increasing our understanding of the oat crown rust pathosystem.

## Acknowledgments

We acknowledge the Australian Grains Genebank, Bruce Winter at the Department of Agriculture and Fisheries (Queensland), and the GRDC-funded oat phenology project team (grant CSP2007) for contributing oat germplasm. We also thank Ben Trevaskis (CSIRO) for helping with pedigree information of several oat cultivars. We acknowledge Liza Apps for technical support at the CSIRO Black Mountain Laboratories Quarantine facility.

## Author contributions

This study was planned and designed by MF and PND. SFK contributed the US oat differential set. ECH, DL, and MF manage collections of rust isolates, virulence assessments, and seed amplification. SB provided DNA material for a subset of oat lines. DTN performed the genotypic analysis. RM assisted with marker design. MM, RS, and JS supported bioinformatics. DTN, ECH, and MF wrote the first draft of the manuscript. All authors contributed to the interpretation of results, reviewed the manuscript, and approved the final version.

