## Supplementary material for "Genotypic and resistance profile analysis of two oat crown rust differential sets urge coordination and standardisation": Figure S

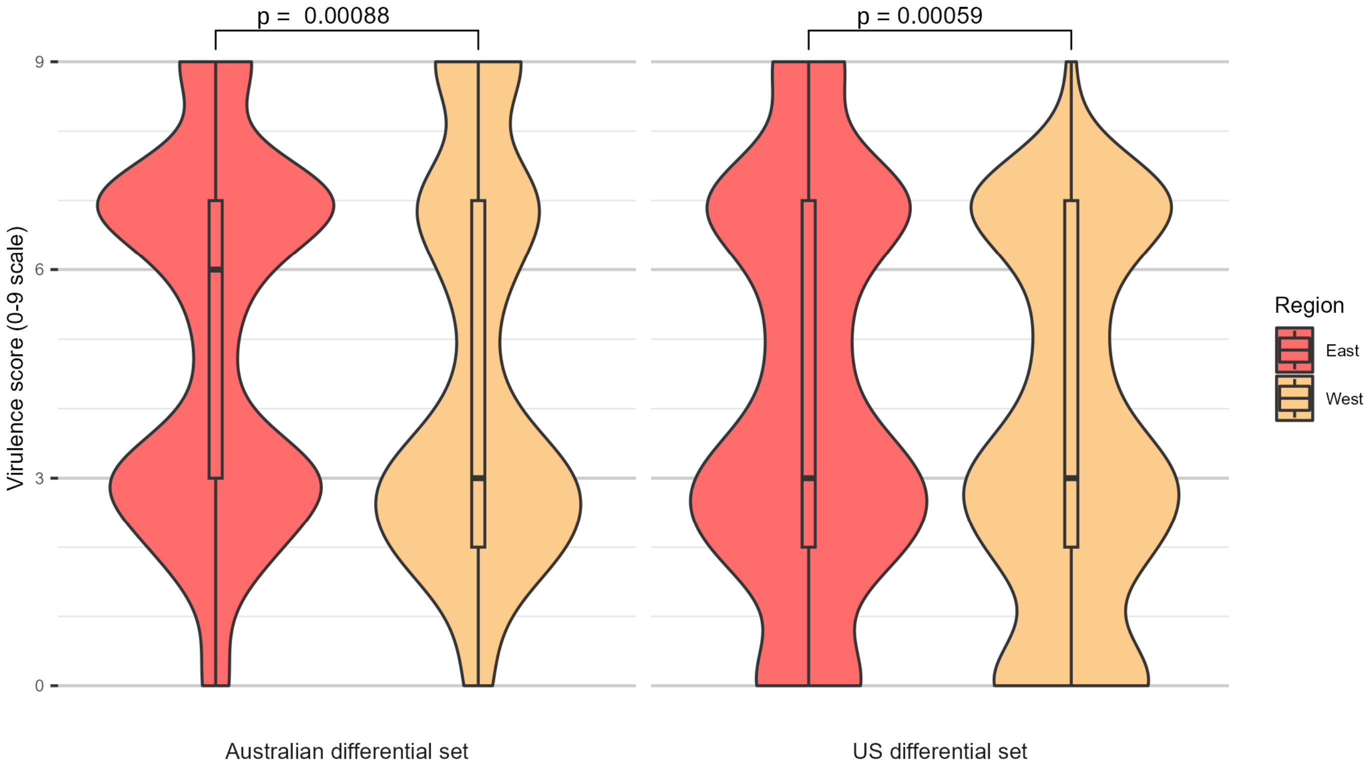


**Figure S1:** Violin plots comparing the pooled virulence scores of 25 *Pca* isolates from the Eastern (red) and Western (yellow) regions of Australia on the Australian and US differential sets. The eastern region includes QLD, NSW, VIC, and SA, and west refers to WA. *P*-values shown are from a two-sided Wilcoxon rank sum test between the virulence scores of the eastern and western isolates on the two differential sets.

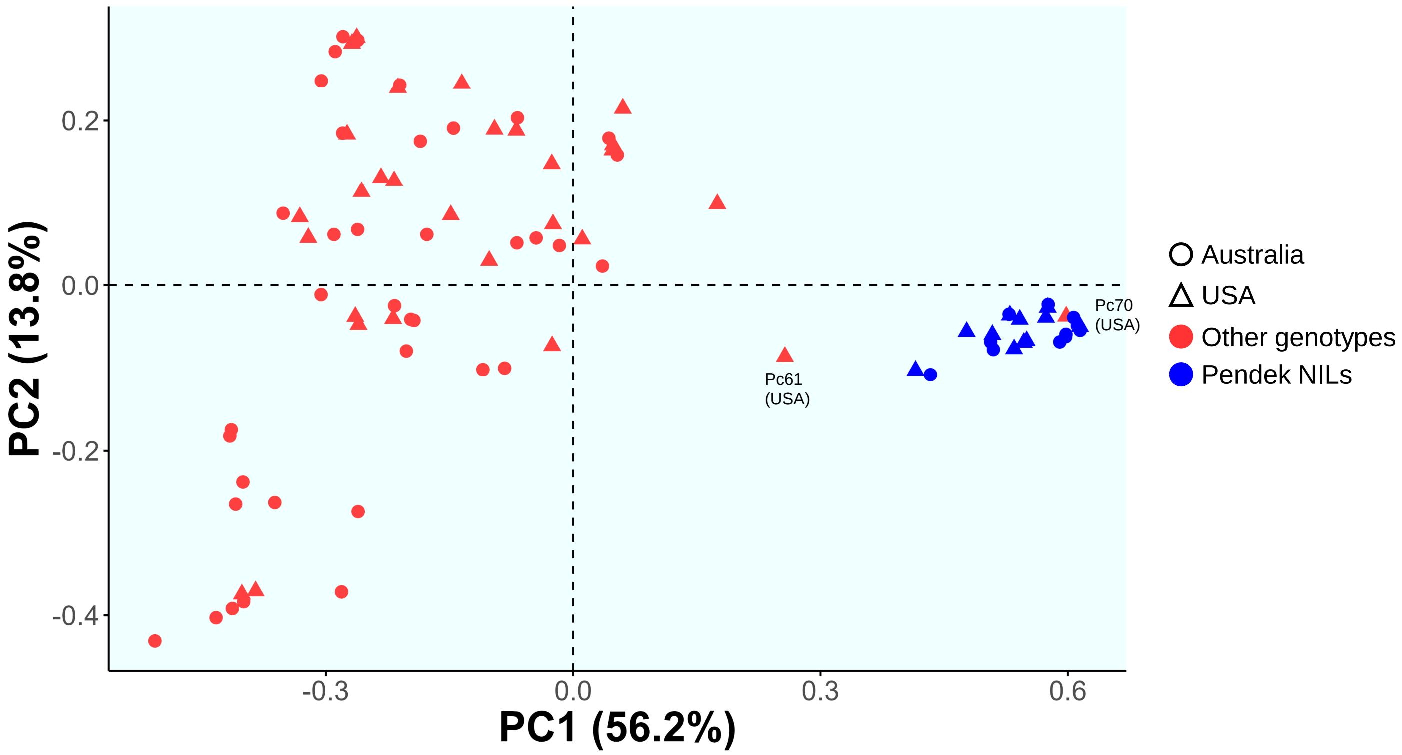


**Figure S2.** Principal component analysis of the oat lines from the Australia and USA differential sets. Dots and triangles indicate oat differential lines representing the USA and Australian sets, respectively. Blue colour indicates lines derived from backcrossing of resistance sources to the oat line Pendek (NILs).

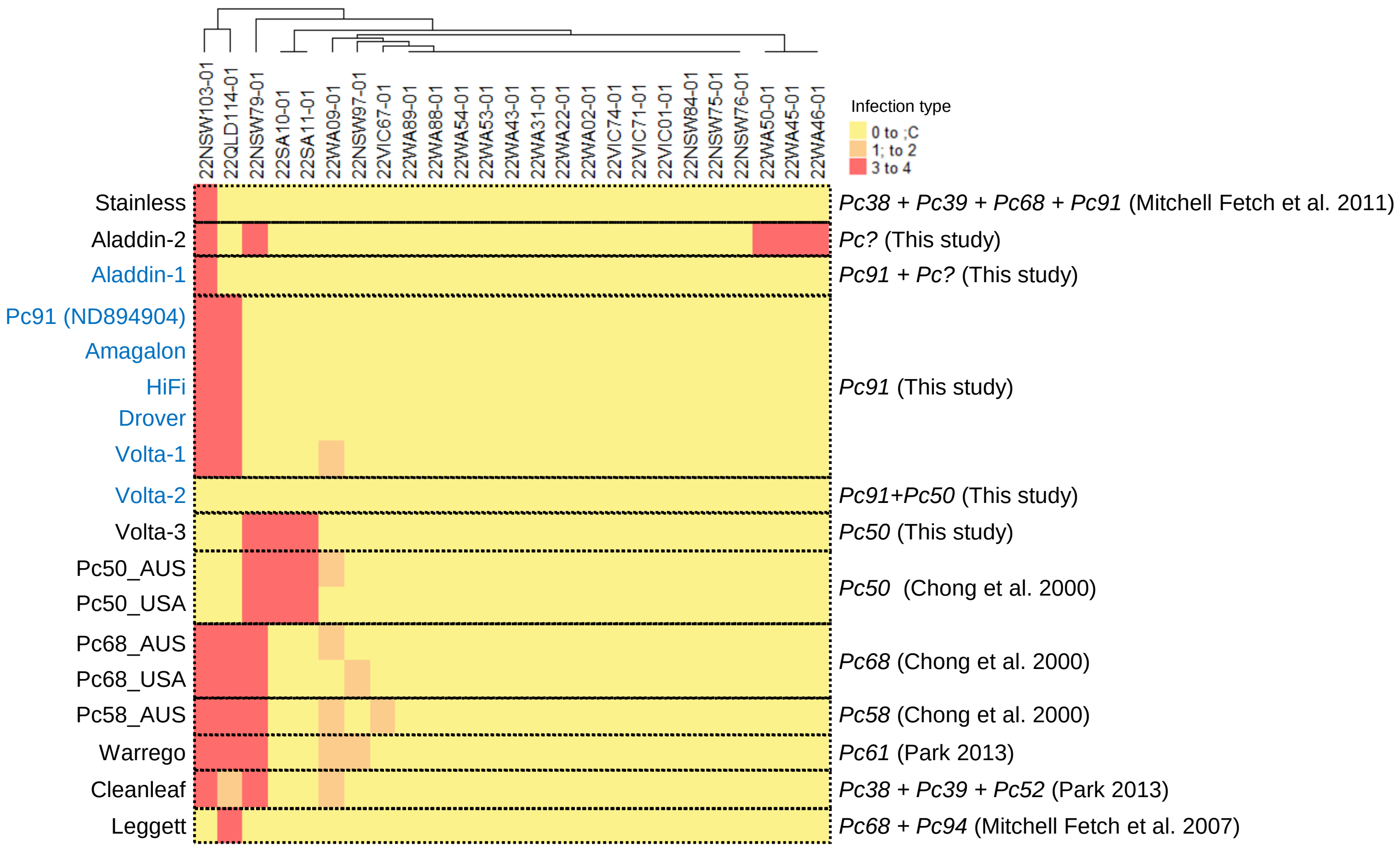


**Figure S3.** Heatmap shows virulence profiles of 25 isolates on *Pc91* carriers and non-carriers oat genotypes. Colour range indicates the infection type of isolate on the host: low virulence (resistance) in yellow and orange (0 to ;C and 1; to 2) and high virulence (susceptibility) in red (3 to 4). Oat lines in blue font are positive for *Pc91* marker *oPt-0350-KOM4c2*. Postulated *Pc* genes based on KASP results, virulence profiles, and previous reports are shown on the right. Aladdin-1, Volta-1 and Volta-2 were derived from AGG seed source (F6-F7), while Aladdin-2 and Volta-3 were derived from a breeder's seed source (F15+).

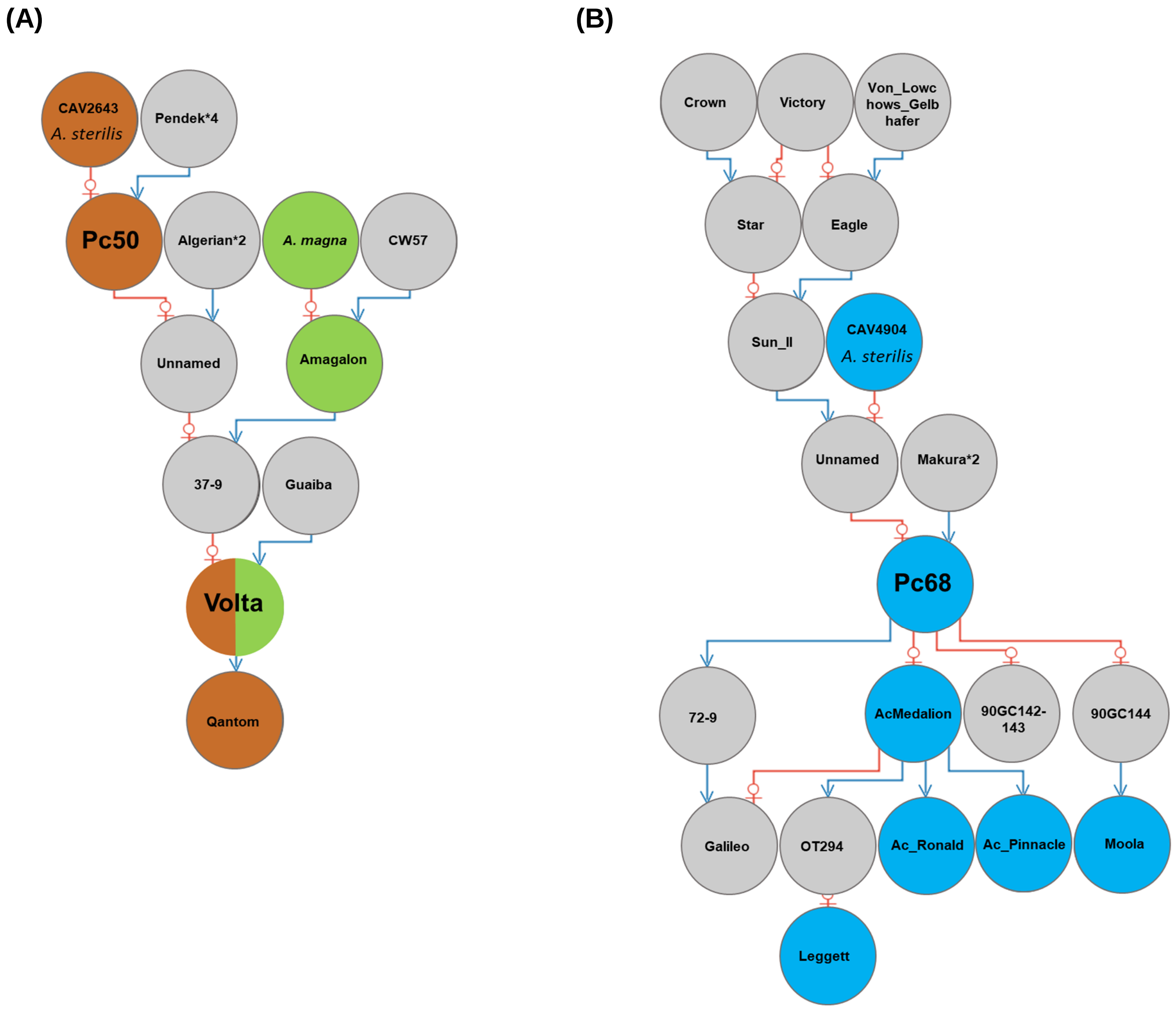


**Figure S4.** Pedigrees involving the oat differential lines **(A)** Pc50 and **(B)** Pc68. Red lines indicate the maternal parent connection and blue indicates the paternal parent. The coloured circles are lines postulated to carry *Pc* genes based on previous historic reports (Brown: *Pc50*, Green: *Pc91*, Blue: *Pc68*).
